# HistoGWAS: An AI Framework for Automated and Interpretable Genetic Analysis of Tissue Phenotypes

**DOI:** 10.1101/2024.06.09.597752

**Authors:** Shubham Chaudhary, Almut Voigts, Michael Bereket, Matthew L. Albert, Kristina Schwamborn, Eleftheria Zeggini, Francesco Paolo Casale

**Affiliations:** Institute of AI for Health, Helmholtz Zentrum München – German Research Center for Environmental Health, Neuherberg, Germany; Helmholtz Pioneer Campus, Helmholtz Zentrum München – German Research Center for Environmental Health, Neuherberg, Germany; School of Computation, Information and Technology, Technical University of Munich, Garching, Germany; TUM School of Medicine and Health, Technical University of Munich and Klinikum Rechts der Isar, Munich, Germany; Department of Computer Science, Stanford University, Stanford, CA, US; Octant Biosciences, San Francisco, CA, US; Institute of Pathology, TUM School of Medicine and Health, Technical University of Munich, Munich, Germany; Institute of Translational Genomics, Helmholtz Zentrum München – German Research Center for Environmental Health, Neuherberg, Germany

## Abstract

Understanding how genetic variation affects tissue structure and function is crucial for deciphering disease mechanisms, yet comprehensive methods for genetic analysis of tissue histology are lacking. We address this gap with HistoGWAS, a framework integrating AI tools for representation learning and image generation with fast variance component models to enable scalable and interpretable genome-wide association studies of histological traits. HistoGWAS employs histology foundation models for automated trait characterization and generative models to visually interpret the genetic influences on these traits. Applied to eleven tissue types from the GTEx cohort, HistoGWAS identifies four genome-wide significant loci, which we linked to specific tissue histological and gene expression changes. A power analysis confirms the effectiveness of HistoGWAS in analyses of large-scale histological data, underscoring its potential to transform imaging genetic studies.

## Introduction

Genetic analysis of quantitative traits has elucidated the functional mechanisms of disease loci identified through genome-wide association studies (GWAS) by pinpointing intermediate processes in disease development. Initially focused on molecular traits such as gene expression and protein levels ^1–7^, these analyses have expanded to include medical imaging-derived traits, enhancing our understanding of intermediate disease processes and uncovering new imaging biomarkers ^8–15^.

Histological images capture a wide range of tissue and cellular phenotypes, offering the potential to study biological processes in disease progression. However, their utility in genetic studies has been limited, primarily due to their vast size and complexity, along with manual and qualitative scoring of disease-related traits, which result in low statistical power and significant challenges in interpretation ^16^.

Recent advancements in artificial intelligence have transformed the way we analyze and interpret complex biological data. Foundation models, for instance, have enabled the derivation of quantitative embeddings from these complex modalities, facilitating various downstream tasks ^17–21^. Concurrently, generative models have improved our ability to study the influences of specific covariates on imaging and molecular traits ^22–24^. However, in the field of computational pathology, despite the significant contributions of AI to diagnostics and prognostics ^25–30^, frameworks harnessing AI for effective genetic analysis of tissue histology are still lacking.

Here, we introduce HistoGWAS, a new AI-enabled framework to enable automated large-scale GWAS of histological traits. Our approach begins with a new semantic autoencoding strategy, utilizing pre-trained foundation models for deriving quantitative histology embeddings for genetic analysis, complemented by generative models to reverse this encoding process (**Figure 1a**). Next, we perform scalable GWAS of the derived embeddings through a novel efficient variance component testing strategy (**Figure 1b**). Finally, we utilize the trained generative model to visualize traits linked to significant genetic loci (**Figure 1c**).

**Figure 1.**
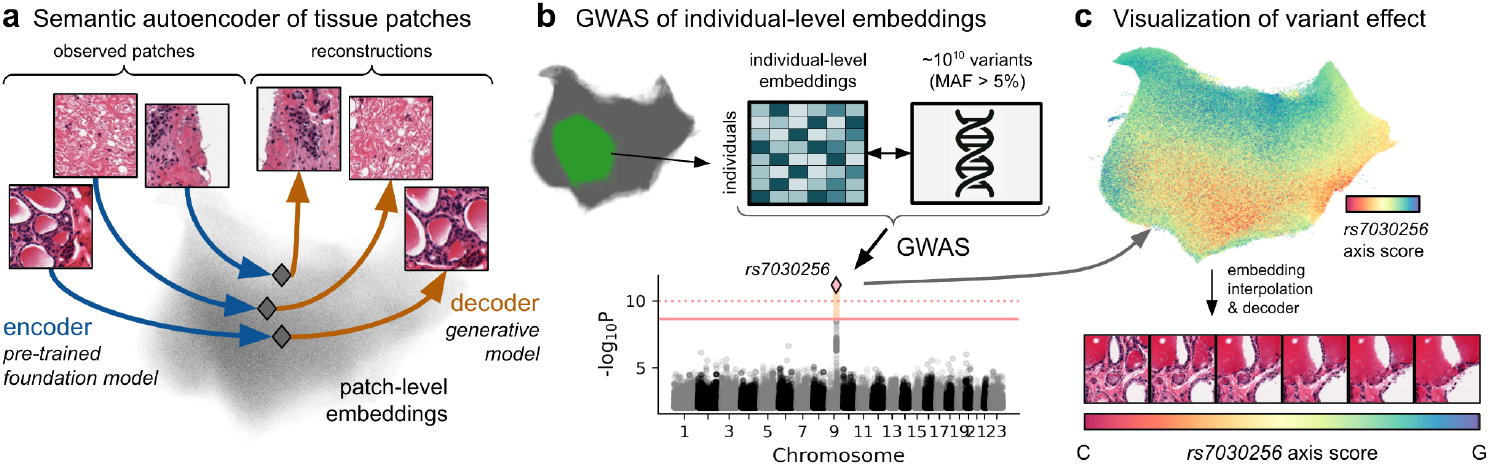
Overview of the HistoGWAS Workflow. (**a**) A semantic autoencoder encodes slide patches into patch-level embeddings and reconstructs these embeddings back into the image space. We utilize a pre-trained foundation model as the encoder and a generative model as the decoder. (**b**) Illustration of a genome-wide association study (GWAS) with HistoGWAS. Patch-level embeddings from similar patches are averaged within each individual to derive individual-level embeddings, which are then associated with genome-wide variants. The output of this analysis is illustrated as a Manhattan plot, showing the P values for association between the individual-level embeddings and single genetic variants. (**c**) Visualization of the histological changes associated with significant variants (e.g., *rs7030256*) is achieved by projecting embeddings —interpolated along the direction of the genetic effect— back into the image space using the semantic decoder.

Notably, HistoGWAS enables, for the first time, the detection of genome-wide significant loci in an analysis of eleven tissues from the GTEx cohort. Furthermore, our simulations demonstrate significant increases in detection power with larger sample sizes, emphasizing the transformative potential of our framework for future imaging genetics studies.

## Results

### Histology Semantic Autoencoders for Joint Analysis with Genetic Data

First, we set out to develop an optimized autoencoding strategy for molecular analysis by identifying the most suitable encoder for molecular predictions and building a decoder capable of accurately inverting this process.

We analyzed histology samples from the Genotype-Tissue Expression (GTEx) dataset ^2^, focusing on eleven tissues with the highest availability of both histology and genetic data (n ≥ 750; **Supplementary Dataset 1**). Following previous studies ^18,31^, we extracted 192μm × 192μm tissue patches from each whole slide image (**Figure 2a**), resulting in a dataset of 19,901,526 patches across 9,006 slides after rigorous quality control (**Supplementary Dataset 1, Methods**).

**Figure 2.**
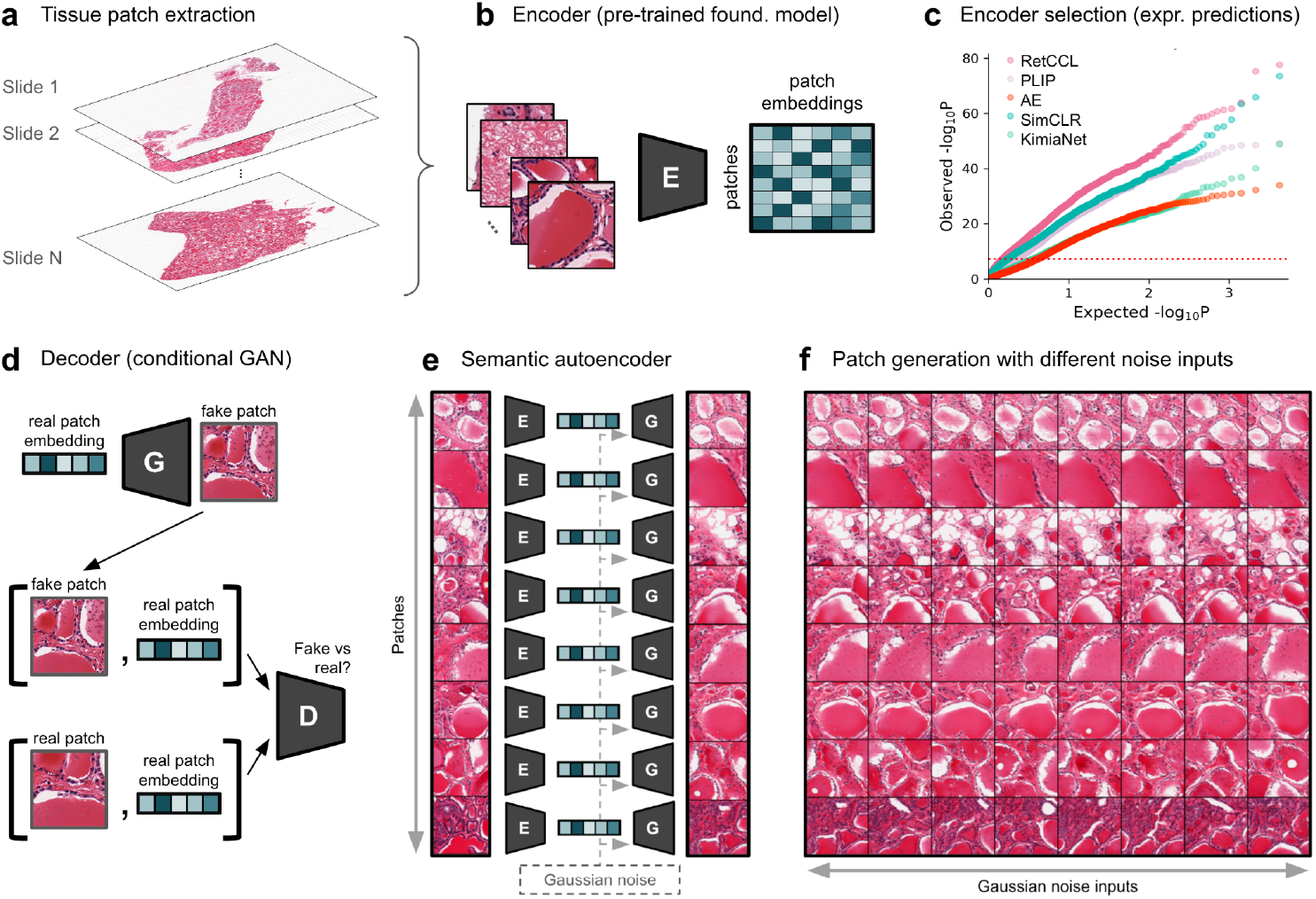
Effective Semantic Autoencoding of Histology Data Using the GTEx Dataset. (**a**) Tissue patches of 192μm × 192μm are extracted from 9,006 whole slide images, totaling 19,901,526 patches after quality control measures (**Methods, Supplementary Dataset 1**). (**b**) Patch-level embeddings are derived using a variety of pre-trained models as semantic encoders: The RetCCL contrastive learning model ^17^, alongside other models such as standard autoencoders ^27,29^, simple contrastive learning methods (SimCLR) ^18,32^, a tissue type classification model (KimiaNet) ^33^, and a pathology foundation model (PLIP) ^17^. (**c**) Comparison of models in predicting gene expression is showcased through a QQ plot of P values of association between observed and predicted expression levels across genes (**Methods**). The best model in this comparison is selected as our semantic encoder. (**d**) A conditional generative adversarial network (GAN) setup trains a generator to produce realistic ‘fake’ patches from embeddings and Gaussian noise, while a discriminator learns to differentiate between real and generated images based on their embeddings, ensuring the generation of high-fidelity reconstructions. (**e**) Encoded patch embeddings and Gaussian noise are fed into the GAN generator, functioning as a decoder, which recreates semantically similar patches. (**f**) Variability in patch generation is shown across different patches with varying Gaussian noise inputs, highlighting the generator’s ability to maintain critical features of the original images.

Starting with the encoder, we determined which of the available pre-trained models is most suited to encode the defined patches into embeddings for genomic analyses (**Figure 2b**). To do so, we assessed how effectively the embeddings from different pre-trained vision models can linearly predict gene expression labels (**Methods**). This comparison encompassed self-supervised contrastive learning methods ^18,32^, supervised models for cancer type classification ^28,33^, a histology foundation model trained across images and pathologist descriptions ^17^, and standard autoencoders directly optimizing image reconstruction ^27,29^. Our findings indicate that contrastive learning methods perform best with the RetCCL contrastive model outperforming all others (average Spearman correlation across genes between predicted and observed values of ρ=0.366±0.008, compared to ρ=0.346±0.008 for simple contrastive learning, ρ=0.310±0.008 for cancer classification models, ρ=0.270±0.010 for the cross-modal foundation model, and ρ=0.230±0.007 for autoencoders; **Figure 2c**; **Supplementary Figure 1**). We thus selected RetCCL as HistoGWAS encoder.

After selecting our encoder, we aimed to develop a decoder that was capable of reverting patch embeddings back to full resolution images. To this end, we trained conditional generative adversarial networks (GANs) ^34–36^ for each tissue type, conditioning patch image generation on the encoder patch embeddings. Briefly, our GAN framework incorporated a generator network tasked with producing ‘fake’ patches from patch embeddings and Gaussian noise, paired with a discriminator network trained to distinguish between real and generated images, based on their corresponding patch embeddings (**Figure 2d, Methods**). After adversarial training, the generator was able to generate realistic images from embeddings, serving as an effective decoder (**Figure 2e**). Semantic reconstructions from this decoder are stochastic, with varying Gaussian noise inputs and fixed embeddings resulting in stochastic reconstructions retaining key features of the original image (**Figure 2f**).

To quantitatively evaluate the quality of our semantic reconstructions, we assessed the ability to predict gene expression from embeddings of actual vs reconstructed images, where reconstructions were obtained either using HistoGWAS semantic autoencoder or a conventional autoencoder. The correlation coefficient (R^2^) for prediction accuracy across genes underscored the superior retention of gene expression signals by our semantic autoencoder, with an R^2^ of 0.653, significantly surpassing the conventional autoencoder’s R^2^ of −0.183 (**Methods, Supplementary Figure 2**).

In summary, we introduced a novel semantic autoencoding strategy that selects an encoder to generate embeddings predictive of molecular features and trains a decoder to invert this process, and demonstrated its superior performance over traditional reconstruction-based autoencoders used in prior histology analyzes ^27,29^.

### Comprehensive Genetic Association Studies across 11 Tissues Reveal Four Genome-wide Significant Loci

To harness the detailed information available in histological slides and effectively capture the subtle genetic influences that may affect specific subregions within tissues, we focused our genetic analyses on 68 distinct cluster signatures identified through a refined clustering analysis within each tissue (**Methods, Supplementary Figure 3, Supplementary Dataset 1**).

**Figure 3.**
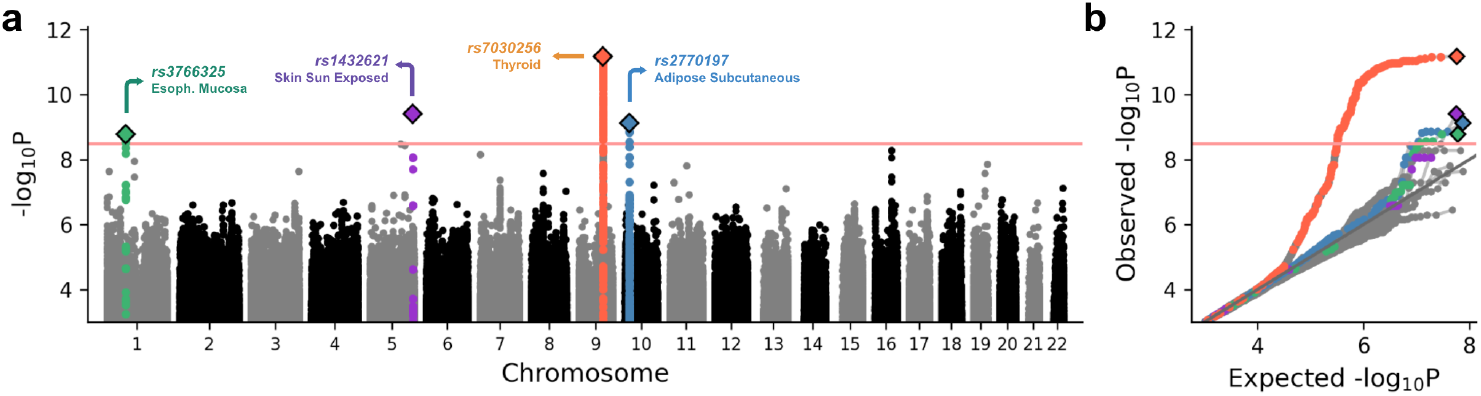
GWAS Analysis of Histological Embeddings Across 68 Cluster Signatures. (**a**) Manhattan plot showcasing the P values for genome-wide association studies (GWAS) of 68 cluster signatures. The red horizontal lines represent thresholds for multi-trait genome-wide significance, set at family-wise error rates of 20%, determined through a permutation-based procedure (**Methods**). Each genome-wide significant locus is marked with a diamond, while variants in linkage disequilibrium are color-coded to match their lead variant (R^2^>0.5). (**b**) QQ plots displaying the P values from genetic association tests, organized by tissue type. Variants are color-coded consistently with (**a**).

First, to validate the biological significance and distinctiveness of these cluster signatures, we conducted correlation analyses between the proportion of patches in each cluster signature and the expression levels of specific genes. This approach uncovered distinct genes and pathways corresponding to different cluster signatures within the same tissue type, revealing the diverse biological functions these signatures represent (**Supplementary Figure 4, Supplementary Dataset 2, Methods**).

**Figure 4.**
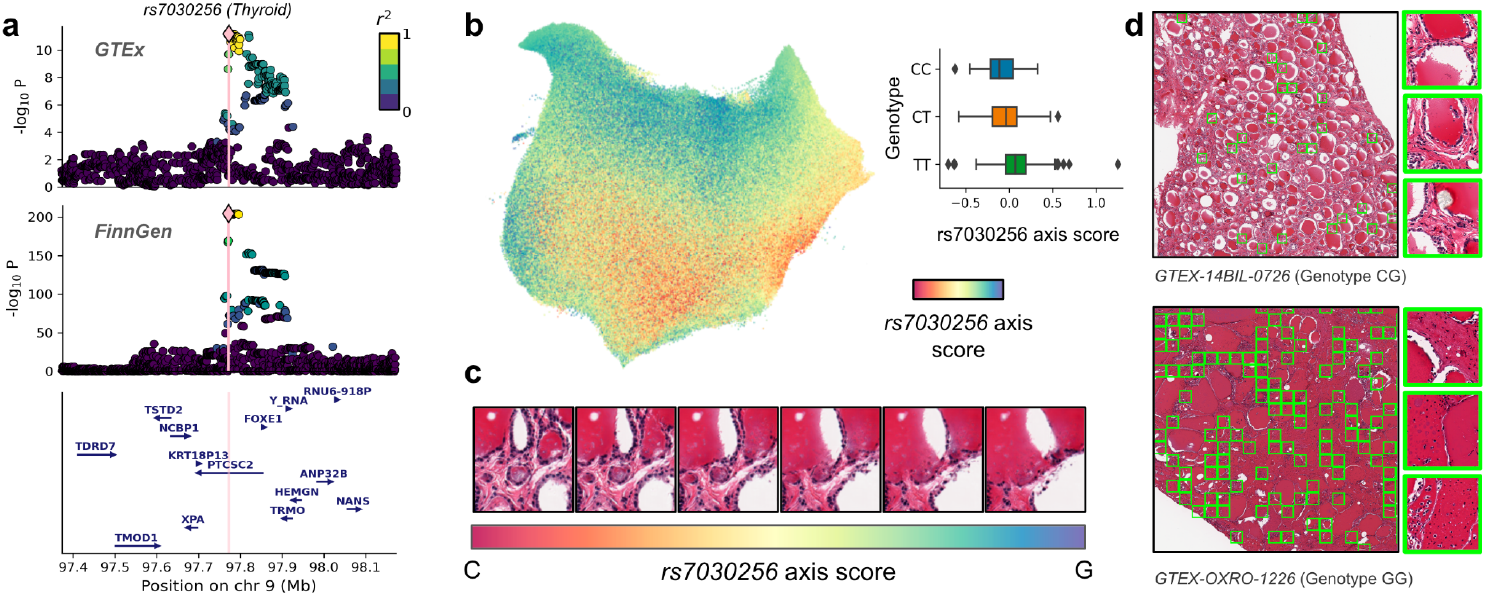
Effect of Intronic Variant *rs7030256* on Thyroid Histology. (**a**) Dual LocusZoom plots illustrating the colocalization of signals for *rs7030256*: the upper plot is derived from GTEx Thyroid Cluster Signature 2, while the lower plot presents findings from the FinnGen GWAS of strict autoimmune hypothyroidism ^40^. Nearby genes are also shown to highlight the genomic context of *rs7030256*. (**b**) UMAP of patch embeddings in thyroid, colored by the inferred *rs7030256* axis scores (**Methods**). An associated box plot displays the variation in these scores across genotypes. (**c**) Histological images demonstrating allele effects by projecting interpolated embeddings —along the direction of the *rs7030256* axis— back into the image space using the semantic decoder. (**d**) Whole slide images showcasing phenotypes associated with different alleles of *rs7030256*. Patches closely linked to specific alleles (defined by the top/bottom 5% percentile of the *rs7030256* axis score) are highlighted in green. For each allele, three patches are magnified to provide a detailed view of the histological differences.

Next, we employed adapted scalable variance component models to enable genome-wide association testing between individual-level average patch embeddings for each cluster signature and approximately 5 million genetic variants (MAF≥5%), while accounting for patient covariates and population structure (**Methods**). Across 68 cluster signatures, this analysis identified four genome-wide significant loci (P < 3.23×10^−9^; corresponding to 20% FWER), accounting for multiple hypothesis testing correction across variants and cluster signatures using a permutation-based procedure (**Figure 3a-b, Supplementary Figure 5, Supplementary Dataset 3, Methods**). Notably, the same analysis on permuted genotypes yielded calibrated P values with no significant associations (**Methods, Supplementary Figure 6**), demonstrating the statistical calibration of the proposed procedure.

**Figure 5.**
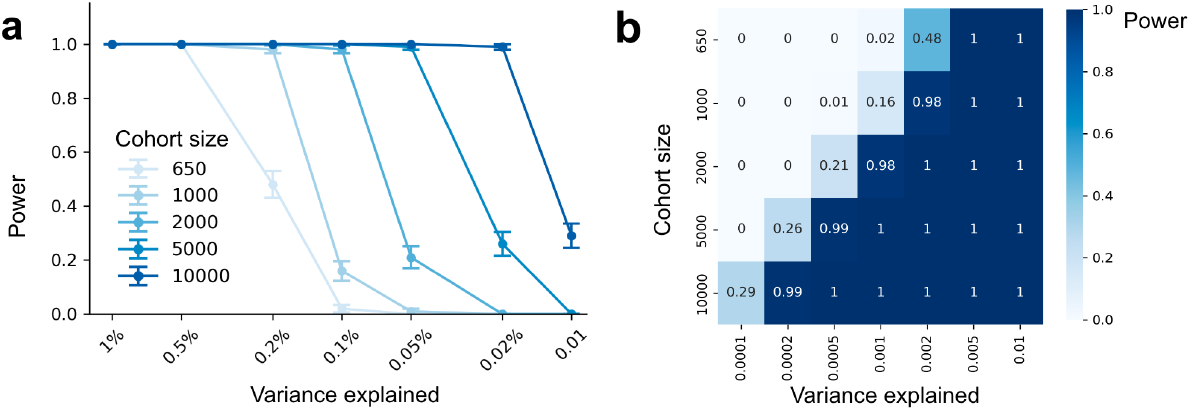
Power Analysis of HistoGWAS Across Varying Sample Sizes and Genetic Effect Sizes. (**a**) Average statistical power of HistoGWAS for identifying single genetic variants, varying the variance explained by the genetic variant and the cohort size. Standard errors are derived from 100 simulation seeds. (**b**) Heatmap illustrating the average statistical power at various combinations of variance explained (rows) and cohort size (columns).

The most significantly associated locus, lead intron variant *rs7030256* in the *PTCSC2* gene (P<6.62×10^−12^), was associated with five different signature clusters in thyroid tissue (**Supplementary Figure 5**). Additionally, three significant loci were each associated with a single cluster signatures in sun exposed skin (*rs1432621*; P<3.94×10^−10^), adipose subcutaneous (*rs2770197*; P<7.39×10^−10^), and esophagus mucosa (*rs3766325*; P<1.64×10^−9^), respectively.

### Biological Characterization of Recovered Genetic Effects on Tissue Histology

To visualize the histological changes linked to each of the identified genetic loci, we utilized our semantic decoder. Briefly, employing the linear mixed model used for association testing, we determined the direction of the genetic effect for each variant in the patch embedding space, termed the ‘genetic effect axis’. We then interpolated along this axis and decoded the resulting embeddings back into the image space using our semantic decoder, which illustrated a continuum of histological changes driven by the genetic variants (**Methods**). Additionally, to probe the biological implications of each locus, we analyzed their effects on gene expression levels, investigated pathways enriched in top associated genes, and examined previously reported associations of each locus with global traits and diseases (**Methods**).

The top variant *rs7030256* of the locus identified in thyroid tissue is an intron variant in the Papillary Thyroid Cancer Susceptibility Candidate 2 gene (*PTCSC2*, **Figure 4a**), a thyroid-specific lncRNA gene associated with thyroid health ^37^. Notably, this signal colocalizes with genetic variants previously identified in GWAS of hypothyroidism in the FinnGen dataset ^38–40^ (**Figure 4a**). Histological assessment of images generated through our interpolation procedure revealed an increase in colloid and enlarged follicles associated with allele G along the *rs7030256* axis (**Figure 4b-c**), indicative of potential goiter development. Furthermore, inflammatory cells were found within the colloid. Conversely, allele C was characterized by a white frame around the colloid, suggesting increased resorption and thyroid activity (**Figure 4b-c; Supplementary Figure 7**). These changes were confirmed in real whole-slide images at lower resolutions (**Figure 4d**). Notably, the *rs7030256* locus also showed associations with changes in the expression levels of 128 genes (Bonferroni-adjusted P < 0.05; **Methods**; **Supplementary Dataset 3**), substantiating its relevance to thyroid physiology. Among the associated genes, we identify *DIO1*, a known regulator of the T3/T4 ratio and hormone secretion ^41^, genes implicated in hypothyroidism such as *GLIS3-AS1* and *DUOXA2* ^42,43^, and several genes linked to Papillary Thyroid Cancer, including *LINC01384, GREB1L, LINC01315, LMO3*, and *MUC1* ^44–48^. Pathway enrichment analysis highlighted hedgehog signaling and the estrogen response as key pathways among associated genes (**Methods**; **Supplementary Dataset 3**), both known for their impact on thyroid cancer progression ^49,50^.

In the tissue from the esophagus mucosa, the lead variant of the detected locus is *rs3766325*, an intron variant in the Interferon-Induced Protein 44 gene (*IFI44*), recognized as an immune biomarker in head and neck squamous cell carcinoma ^51^ and specific infectious diseases ^52^. Histologically, allele A of *rs3766325* was associated with notable enlargement of epithelial cell nuclei in the mucosal layer of the esophagus (**Supplementary Figure 7, Supplementary Figure 8**), indicative of post-injury regeneration ^53^. Gene expression analysis identified 30 genes associated with the *rs3766325* genotype (Bonferroni-adjusted P < 0.05, **Methods**). Genes upregulated with allele A, such as *LAPTM4B* and *MELK*, were linked to enhanced cell proliferation, whereas downregulated genes, including *MAL* and *CRCT1*, inhibited this process ^54–57^, which is in line with the observed histological differences. Finally, pathway enrichment analysis was also consistent with these observations, highlighting upregulation of *E2F* target ^58^ and *MYC* transcription factor targets pathways ^59^, and downregulation of the p53 tumor suppressor gene ^60^ with allele A (**Supplementary Dataset 3**).

For lower leg tissue from sun-exposed skin, the intergenic lead variant *rs1432621* on chromosome 5 was histologically characterized by allele G association with increased collagen density in the dense irregular connective tissue (**Supplementary Figure 7, Supplementary Figure 8**). Although no single gene expression was significantly associated with the *rs1432621* genotype after multiple hypothesis testing correction, pathway enrichment analysis in top 50 associated genes revealed relevant pathways, including UV response, glycolysis, and epithelial mesenchymal transition mechanisms ^61–63^ (**Supplementary Dataset 3**). In subcutaneous adipose tissue, the allele A of the lead variant *rs2770197* was histologically linked to an increase in vacuole size and cell membrane deterioration, indicators of tissue degradation (**Supplementary Figure 7, Supplementary Figure 8)**. Top 50 associated genes enriched for adipogenesis, fatty acid metabolism, estrogen response (Early), and *mTORC1* Signaling, all playing crucial roles in adipocyte appearance and obesity ^16,64,65^ (**Supplementary Dataset 3**).

### Power Analysis Demonstrates HistoGWAS’ Effectiveness and Scalability

To gauge HistoGWAS’ efficacy for advancing histogenetics studies, we undertook a power analysis, simulating cohorts with up to 10,000 individuals. Briefly, we simulated trait embeddings with influences from covariates, a genetic variant and Gaussian noise (**Methods**). After confirming calibration of P values under the null (no simulated genetic effects) (**Supplementary Figure 9**), we assessed statistical power of detecting the simulated causal variant with HistoGWAS at P<5×10^−8^, for varying sample sizes and genetic effect sizes.

In scenarios similar to our study’s scale (650-1000 individuals), we had high power (>95%) for variants explaining at least 0.2% of the variance in the embeddings space (**Figure 5a**). Notably, as we increased sample size, the ability to detect subtler genetic effects significantly improved: we could detect variants that explain ≥0.1%, ≥0.05% and ≥0.02% with 2,000, 5,000 and 10,000 individuals, respectively. The heatmap in **Figure 5b** shows the sample sizes needed to achieve the desired statistical power to detect genetic effects with a given magnitude, serving as a guide for the design of future histogenetic cohorts.

## Discussion

We introduce HistoGWAS, an integrated AI-enabled framework for automated and scalable genetic analysis of tissue phenotypes leveraging large histology cohorts. This is achieved through the integration of a novel semantic autoencoder for semantic compression of tissue phenotypes with scalable variance component models enabling association testing between single genetic variants and the derived imaging representations. This setup allows for the visualization of genetic effects on tissue structures and enables the examination of tissue changes mediating the effect of genetic loci on disease risk, thereby extending genetic analyses of intermediate phenotypes to tissue histology for the first time ^1–7^.

Applying HistoGWAS to data from eleven tissue types within the GTEx cohort yielded four significant loci (P<2.29×10^−9^). A particularly noteworthy finding is the identification of the *PTCSC2* locus in thyroid tissue (P<6.62e^-12^), a thyroid-specific lncRNA gene linked to thyroid health ^37^. The top variant *rs7030256* of the locus is characterized by pronounced histological changes including the increase in colloid and enlargement of follicles, consistent with non-toxic multinodular goiter. The locus colocalizes with a previously detected genetic signal for hypothyroidism and general thyroid issues ^40^, which demonstrates the ability of HistoGWAS to uncover disease mechanisms by leveraging genetic and histological data.

Moreover, while the GTEx cohort’s unique composition poses challenges for independent validation, power analysis confirms that HistoGWAS is exceptionally effective in large-scale histology cohorts, demonstrating its potential to pioneer the new field of histo-genetic analyses. Here, focusing on germline genetic effects, we enable the search of causal histology features to understand disease, distinguishing our analysis from previous descriptive approaches ^27–30^. When considering this alongside the generality and scalability of the framework—utilizing a pretrained foundation model, simple decoding, and scalable association testing—HistoGWAS is poised to become the gold standard for detailed characterization of genetic effects on tissue structure and function and their role in disease progression.

Finally, the approach of integrating advanced AI tools for the encoding and decoding high-content phenotypes with multivariate statistical analysis of latent spaces is applicable to a wide array of high-content phenotypic modalities and use cases. For example, the approach implemented in HistoGWAS can facilitate the study of genetic effects on morphological changes visible through non-invasive medical imaging ^8–14^, and enable the assessment of the impacts of genetic and chemical perturbations in high-content screens of in vitro systems ^66^. This is particularly relevant in advanced systems like organoids, enabling a detailed description and visualization of complex phenotypic changes resulting from induced perturbations.

## Methods

### Preprocessing and Encoding of Tissue Histology Data

#### Data Preprocessing and Patch Extraction

An initial dataset was curated from the GTEx project, focusing on tissues for which both histological and genetic data was available for a minimum of 750 individuals. This resulted in the inclusion of 11 tissue types, encompassing a total of 9,006 slides (**Supplementary Dataset 1**). We delineated patches measuring 192μm × 192μm for each analyzed slide, employing a pre-established grid. The process involved converting low-resolution slides to grayscale, followed by the application of the binary thresholding *cv2*.*threshold* function from the Opencv library ^67^, to differentiate between tissue (foreground) and the absence of tissue (background). This mask was then utilized to select patches containing at least 50% tissue, which were exported as 256 × 256 images (0.75 μm per pixel). This process resulted in a dataset containing 19,901,526 patches from 11 different tissues (see **Supplementary Dataset 1**).

#### Compared Models for Semantic Encoding

We evaluated embeddings from four distinct pretrained models as candidate semantic encoders for HistoGWAS: (i) RetCCL ^17^, pretrained on the TCGA dataset using cluster-guided contrastive learning, (ii) SimCLR^18,32^, pretrained on the ImageNet dataset employing a standard contrastive learning procedure, (iii) KimiaNet ^33^, pretrained on TGCA for cancer type classification, (iv) PLIP ^17^, pretrained on slide images and associated pathologist descriptions from OpenPath through a multimodal contrastive-learning framework. All models were applied to the GTEx dataset without fine tuning. In addition to pretrained models, we trained a classical autoencoder (AE) on GTEx for each of the 11 tissues, which aimed to optimize reconstruction using an L2 reconstruction loss (**Supplementary Information**). This approach has been considered in previous analyses of histology and genomics dataset in the GTEx dataset ^27,29^. After defining embeddings for all foreground patches, we conducted Principal Component Analysis (PCA) within each tissue, and reduced embeddings to their leading 64 principal components. This decision was informed by a simulation analysis that evaluated the calibration of P values in genome-wide studies using HistoGWAS under a null model (**Supplementary Figure 10**; see also the power analysis section).

#### Encoder Performance Evaluation and Selection

Across the five compared models, we aimed to select the model whose embeddings could best predict gene expression levels across the 11 analyzed tissues. Briefly, for each tissue, individual-level embeddings were computed as the mean of patch-level embeddings from the same individual. We then employed a variance component model for each gene, with gene expression as the outcome, and effects from individual-level embedding as random effects. For this analysis, we considered the log_10_ TPM expression values obtained from the GTEx Analysis V8 Open Access Datasets ^68^, targeting highly-variable genes within each tissue, identified using the scanpy *highly_variable_genes* function ^69^. This model was fitted on 50% of the individuals (training set), and then used to make gene expression predictions on the remaining 50% (test set). Prediction accuracy was evaluated using Spearman’s correlation between observed and predicted gene expression levels across individuals in the test set, with significance determined by Bonferroni-corrected Spearman P values < 0.05. Among the evaluated models, RetCCL emerged as the most accurate in gene expression prediction and was consequently chosen for all further analyses with HistoGWAS (**Figure 2c, Supplementary Figure 1**).

### Development of Generative Model for HistoGWAS Decoder

To invert the encoding process, we train a generator within the conditional Generative Adversarial Networks (cGAN) framework ^34^, specifically adapted to condition on patch embeddings from the HistoGWAS encoder. This adaptation enables the generator to create synthetic patch images from observed patch embeddings, while a discriminator learns to differentiate between real and generated images based on those embeddings. For generating high-resolution 256×256 images, we adopted the progressive training strategy from Progressive GANs ^36^, gradually adding new layers to both the generator and discriminator to capture increasingly finer details during the training process ^36^.

#### Architecture of the Generator network

Our model processes a 512-dimensional latent representation *z* through a convolutional function *G*(*z*) to generate an image. While in classic GANs *z* is sampled from a standard normal, we here condition its generation on 64-dimensional patch embeddings *x* by assuming *z* is drawn from a multivariate normal distribution with mean *m*_*z*_ (*x*) and standard deviation *s*_*z*_(*x*), both functions of *x*. By rewriting *z = m*_*z*_(*x*) + *s* _*z*_ (*x*) ⍰ ε, with ε∼*N*(0, 1) and ⍰ denoting the Hadamard product, we can separate the dependency on *x* from the stochasticity and enable backpropagation via the reparametrization trick ^70^. We employ linear layers for both *m* _*z*_(*x*) and *s*_*z*_(*x*), with *s*_*z*_(*x*) using a softplus activation function to ensure a non-negative output. Our implementation adapts the publicly available Progressive GAN architecture from Facebook ^71^. For detailed specifications of the generator architecture, we refer to this code base.

#### Architecture of the Discriminator Module

Similar to ^36^, our discriminator processes a patch image *y*, producing a 512-dimensional intermediate representation *h* through the convolutional function *D*(*y*). Given *h*, the output of the discriminator is scalar for each patch, and is the sum of an unconditional component (a linear layer applied to *h*) and a conditional component. The conditional component is derived from the dot product between *h* and a 512-dimensional encoding of the observed patch embedding, obtained from patch embeddings via a linear layer. This conditioning approach is implemented in BigGAN to condition on per-class learnable encodings ^72^, but we here replace learnable encodings with a learnable linear function of the observed patch embeddings. For detailed specifications of the discriminator architecture, we refer to the Facebook code base ^71^.

#### Progressive training and optimization details

We trained our model using the Wasserstein GAN loss ^73^ with gradient penalty to improve stability and convergence (λ=10). Following ^71^, the training initiates with 4×4 pixel image generation, progressively increasing the resolution by adding layers to both generator and discriminator through stages: 8×8, 16×16, 32×32, 64×64, 128×128, and culminating in 256×256 images. Each resolution phase is designated a set number of iterations — 48,000 for the initial and 96,000 for subsequent stages, where a batch of 64 patch images was processed at each iteration. We employed the Adam optimizer ^74^, with β_1_ set to 0, β_2_ to 0.99, and a learning rate of 0.01, optimizing both networks iteratively to balance the generator and discriminator learning processes ^75^.

#### Validation of Semantic Reconstruction Accuracy

Our semantic autoencoder combines the RetCCL model with top 64 principal component extraction for the encoding process, and the cGAN generator for decoding. After visually assessing original and reconstructed patches under varied noise inputs (**Figure 3c**), we proceeded for a quantitative validation using gene expression. Specifically, we compared the quality of gene expression prediction from embeddings of actual versus reconstructed images. Briefly, for each tissue, we fitted a variance component model on 50% of individuals using individual-level embeddings from real patches as inputs. After training, we used this model to predict gene expression for the remaining 50% of the individuals using as inputs individual-level embeddings from either original or reconstructed patches. Consistency of gene expression prediction accuracy from embeddings of actual images versus reconstructed images was quantified as the coefficient of determination (R^2^) between prediction test statistics across genes—R^2^=1 would indicate full retention of expression signal (**Supplementary Figure 2**). We compared retained expression signals of our semantic autoencoder versus a conventional autoencoder, showcasing a marked improvement (**Supplementary Figure 2**). A full presentation of our evaluation procedure can be found in **Supplementary Information**.

### Genetic Analysis of Histological Embeddings

#### Definition of Histological Cluster Signatures for GWAS

To optimally capture the diverse histological phenotypes within each tissue for genetic analysis, we performed an unsupervised analysis of RetCCL embeddings using the *scanpy* Python module ^69^. Briefly, for each tissue, dimensionality was reduced to the 64 leading principal components via Principal Component Analysis (PCA). We then constructed the nearest neighborhood graph (n_neighbors=10), applied Uniform Manifold Approximation and Projection (UMAP) ^76^, and executed Leiden clustering (resolution=0.5) ^77^. These hyperparameters were optimized to discern large, homogeneous clusters showcasing morphological consistency (**Supplementary Figure 3-4**). To ensure clusters had sufficient representation for downstream genetic analyses and to filter out imaging artifacts and tissue impurities, we retained clusters represented by at least 10 patches per slide in at least 650 distinct slides. This approach yielded 68 distinct tissue signature clusters for subsequent analysis (**Supplementary Figure 3-4**), containing 19,901,526 patches (**Supplementary Dataset 1**).

#### Associating Cluster Signatures with Gene Expression

To validate the histological cluster signatures derived from image analysis, we assessed their correlation with gene expression levels. For each signature, we calculated its abundance within slides—defined as the proportion of patches per signature—and analyzed its association with gene expression using a linear model, with Gaussianized gene expression levels as the dependent variable. P values of association were assessed via a log-likelihood ratio test. QQ plots for each cluster signature, highlighting the top five genes associated with each signature and showcasing examples of histological patches from that signature are displayed in **Supplementary Figure 4**. Full results from this analysis can be found in **Supplementary Dataset 2**.

#### Association testing framework

To assess genetic associations with histological traits, we employed a variance component within a linear mixed model framework. Briefly, given the genotype vector **g** for a variant of interest across *N* individuals, the *N* × *L* matrix of individual-level embeddings **X**, and the *N* × *K* covariate matrix **F** of *K* covariates, we considered the generalized variance component model:

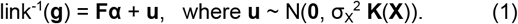

Here, the link function depends on the chosen likelihood for genotype values, **α** denotes the covariate effects, and **K**(**X**) is an *N* × *N* cosine similarity-based covariance function, which describes pairwise similarities between individuals based on their histological embeddings **X**. This model enables association testing by assessing σ_X_^2^ > 0 through a variance component score test ^78^, analogous to SKAT models ^79,80^. We explored both binomial ^81,82^ and Gaussian likelihoods for genotype values, opting for the latter due to its computational efficiency and statistical calibration—despite potential P value inflation due to model mismatches. The embedding dimension *L* represents the number of principal components on RetCCL embeddings, performed separately within each tissue. To select *L*, we evaluated the calibration of our statistical tests under the null model through simulations (**Supplementary Information**), which indicated *L =* 64 as a reasonable compromise due to increasingly deflated P values with larger numbers of components (**Supplementary Figure 10**). Within this framework, we tested for association between individual-level embeddings of 68 cluster signatures and approximately 5 million genetic variants with minor allele frequency ≥ 5%, while accounting for sex, age, type of death, and the four leading genetic principal components. More info about the selected covariance function, efficient computations in the testing procedure and the choice of the likelihood model can be found in **Supplementary Information**.

#### Multiple hypothesis testing correction

To account for multiple hypothesis testing across both genetic variants (MAF ≥ 5%) and cluster signatures, while also considering test dependence due to linkage disequilibrium, we employed a permutation-based approach to control the family-wise error rate (FWER). We conducted 100 permutations of the genetic data for each of the 68 cluster signatures, yielding a total of 6,800 genome-wide association analyses under the null hypothesis. By taking the minimum P value across all variants for each permutation, we obtained a distribution of 6,800 minimum P values. We used the 20th percentile of this distribution to empirically determine the 20% FWER threshold (α=0.2). To adjust for multiple testing across the 68 cluster signatures, this threshold was further divided by the number of clusters. This procedure resulted in a stringent P value threshold of P < 3.23×10^−9^.

### Characterization of Genome-wide Significant Genetic Loci

#### Visualization of genetic effects on histology

To illustrate histological alterations linked to significant genetic variants, we implemented a method integrating embedding interpolation with semantic decoding. Initially, we applied a variance component model to establish the direction of each genetic variant’s effect within the embedding space, termed the ‘genetic effect axis’. For the interpolation process, we calculated extreme embeddings from patches at both extremes of the path-level projection scores along the genetic effect axis—specifically, we considered average embeddings across the patches in the 1st-5th and the 95th-99th percentiles of the projection score, respectively. Interpolating between these extreme embeddings and translating them back into image space via our semantic decoder allowed us to visualize a continuum of histological transformations associated with the genetic variant’s influence. Given the stochastic nature of our decoder, multiple visual interpretations of this semantic interpolation can be generated by varying the input noise, enabling thorough evaluation (**Supplementary Figure 11**). A comprehensive description of this procedure can be found in the **Supplementary Information**.

#### Association with expression levels and global traits and diseases

To characterize genome-wide significant loci, we analyzed their influence on gene expression, biological pathways, and broader traits. Gene expression impact was assessed by testing the association between Gaussianized expression levels of highly variable genes in relevant tissues and the lead locus variant, controlling for sex, age, type of death, and four principal genetic components. Associations were deemed significant at a Bonferroni-adjusted P < 0.05. For pathway analysis, we employed Fisher’s exact test using the *enrichr* tool within the *gseapy* Python module ^83^, using the *MSigDB_Hallmark_2020* annotations ^84^, focusing on the top 50 genes positively or negatively associated. The top five pathways with altered activity, irrespective of P values, were selected for further interpretation. Additionally, the open target genetics interface was used to explore the loci’s links with global traits ^85^.

### Power analysis

To assess HistoGWAS’s statistical power under diverse scenarios, we performed a power analysis, simulating individual-level embeddings as additive contributions from simulated covariates (sex, age, and genetic principal components), a genetic variant, and Gaussian noise. Associations between these simulated embeddings and genetic variants were evaluated using the HistoGWAS testing framework and power was assessed at FWER 5%. Comprehensive simulation details are provided in **Supplementary Information**. Power estimations varied across cohort sizes (650, 1,000, 2,000, 5,000, 10,000) and the variance explained by the genetic variant (1%, 0.5%, 0.2%, 0.1%, 0.05%, 0.02%, 0.01%), employing 100 random simulation seeds for each scenario.

## Declarations

### Ethics approval and consent to participate

No new data was generated for this study. The different ethics approval for the GTEx dataset can be found in the corresponding publications ^2^. The research conformed to the principles of the Declaration of Helsinki.

### Declaration of generative AI and AI-assisted technologies in the writing process

During the preparation of this work the authors used the large language model GPT-4 (https://chat.openai.com/) for editing assistance, including language polishing and clarification of text. After using this tool/service, the authors reviewed and edited the content as needed, and take full responsibility for the content of the publication.

## Supporting information

suppInfo.pdf

## Acknowledgements

We would like to thank Thomas Schwarz-Romond for feedback on the manuscript. This research was conducted using data from the GTEx database, under dbGaP application number #32009. F.P.C. and S.C. received funding by the Free State of Bavaria’s Hightech Agenda through the Institute of AI for Health (AIH). Results from the FinnGen study allowed for the linking of the described gene locus in thyroid and various medical conditions.

## Authors’ contributions

S.C., M.B. and F.P.C. developed the methods. S.C. carried out the experiments and data analysis. A.V., M.A., K.S., and E.Z. provided critical insights for interpreting the results. K.S. offered expert guidance to describe histological phenotypes. F.P.C. conceived the study and supervised the work. The initial draft was written by S.C., A.V., and F.P.C., with all authors contributing to subsequent revisions and refinements of the manuscript.

## Availability of data and materials

### Data availability

No primary data were generated for this study. The GTEx v8 dataset is available at https://gtexportal.org/home (under dbGaP protection). All histological slides mentioned in the paper and the figures can be accessed and analyzed under https://www.gtexportal.org/home/histologyPage.

### Code availability

- RetCCL: https://github.com/Xiyue-Wang/RetCCL
- SimCLR: https://lightning-bolts.readthedocs.io/en/0.3.4/models_howto.html
- KimiaNet: https://github.com/KimiaLabMayo/KimiaNet
- PLIP: https://github.com/PathologyFoundation/plip
- Progressive GAN: https://github.com/facebookresearch/pytorch_GAN_zoo

