## Supplementary material for "HistoGWAS: An AI Framework for Automated and Interpretable Genetic Analysis of Tissue Phenotypes": suppInfo.pdf

### Supplementary Information for “HistoGWAS: An AI-enabled Framework for Automated Genetic Analysis of Tissue Phenotypes in Histology Cohorts”

**2 Supplementary Datasets** **8**

**3 Supplementary Figures** **9**

#### 1 Supplementary Methods

##### 1.1 Standard Autoencoder

In our analyses, we compared the HistoGWAS semantic autoencoder with a standard autoencoder optimized for image reconstruction, which we implemented following previous work leveraing the GTEx resource for molecular analyses. We considered the same architecture and optimization procedure as implemeted in [1], optimizing the mean square loss, between the reconstructed adn origianl images using the Adam optimizer with learning rate 1e-4. Full details on its architecture and optimization details can be found below.

**Architecture of encoder.** The standard autoencoder consists of two primary components: encoder and decoder. The encoder consists of a series of five convolutional layers, each followed by a max-pooling layer and ReLU activation. The convolutional layers incrementally increases the number of channels from 3 to 128 (3, 16, 32, 64, 128), maintaining a constant kernel size of 3 and stride of 1 with padding. Following convolutional layers, a flattening step transitions the data from 3D tensor to 1D tensor followed by a linear layer to produce the encoded representation of dimension 1024.

**Architecture of decoder.** The Decoder reconstructs the input image from encoded representation. Starting with a linear layer that expands the representation back to spatial dimension (128 channels and 8×8 spatial size), then a series of upsample operations followed by convolutional layers and ReLU activation excepts. Through these operations the channels incrementally decrease from 128 to 3 (128, 64, 32, 16, 3). We exceptionally use the Tanh activation (rather than ReLu) after the last convolutional layer.

#### 1.2 Procedure for Validating the HistoGWAS Semantic Autoencoder using Expression Data

This section outlines the step-by-step process used to quantitatively validate our semantic autoencoder reconstructions using gene expression data.

1. **Compute Individual-Level Embeddings from Real Patches**
   - Obtain patch-level embeddings from real patch images using encoder (e.g., RetCCL);
   - Train and apply Principal Component Analysis (PCA) on real patches (dimensionality is reduced to 64 PCs based on calibration analysis (**Supplementary Figures 10**);
   - Average these reduced patch-level embeddings within individuals to obtain individual-level embeddings.
2. **Compute Individual-Level Embeddings from Reconstructions**
   - Start from reduced patch-level embeddings obtained from real patches;
   - Sample reconstructions from these embeddings using the decoder;
   - Re-encode these reconstructed patches using the same encoder as in Step 1;
   - Apply the same PCA used in Step 1 to these re-encoded embeddings to reduce dimensionality;
   - Average these reduced patch-level embeddings within individuals to obtain individual-level embeddings.
3. **Train Variance Component Model and evaluate Out-of-Sample Predictions.**
   - For each gene: (i) Train a linear mixed model on 50% of the individuals to predict gene expression from the individual-level embeddings from real patches. The linear mixed model has $\log_{10}$ TPM (Transcripts Per Million) as outcome, a fixed effect from the intercept, and random effects from individual-level embeddings; (ii) Perform out-of-sample predictions of gene

expression on the remaining 50% of the individuals using individual-level embeddings from either real or reconstructed patches as inputs. These can be obtained through the Best Linear Unbiased Predictions (BLUP) [6];

- Compare the prediction accuracy of gene expression from real versus reconstructed patches across genes to evaluate how much of the expression information is preserved in the reconstructions. This is done using the coefficient of determination ($R^2$) between prediction test statistics, $-\log_{10}$(Spearman $P$ value). The value $R^2 = 1$ indicates full retention of gene expression information.

#### 1.3 Association Testing Framework

**Mixed Model for Association Testing.** We employed a linear mixed model framework to assess genetic associations with histological traits. Specifically, for a genotype vector $\boldsymbol{g}$ across $N$ individuals, the $N \times L$ matrix of individual-level embeddings $\boldsymbol{X}$, and the $N \times K$ covariate matrix $\boldsymbol{F}$ of $K$ covariates, we utilized the following generalized linear mixed model:

$$\text{link}^{-1}(\boldsymbol{g}) = \boldsymbol{F}\boldsymbol{\alpha} + \boldsymbol{u}, \quad \text{where}\, \boldsymbol{u} \sim \mathcal{N}(\boldsymbol{0}, \sigma_X^2 \mathcal{K}(\boldsymbol{X})), \tag{1}$$

where $\boldsymbol{\alpha}$ represents the effects of covariates, and $\mathcal{K}(\boldsymbol{X})$ is an $N \times N$ covariance function that models pairwise similarities between individuals based on their histological embeddings $\boldsymbol{X}$. The link function connects the predicted values to the genotype vector $\boldsymbol{g}$, converting the relationship between them.

**Choice of Likelihood.** We evaluated two likelihood functions for this analysis:

1. *Binomial (two trials):* This method, previously used for modeling genotype minor allele counts [7, 5], leverages a binomial distribution with two trials, suitable for genotypes often represented as counts of minor alleles (0, 1, or 2). The logistic sigmoid function acts as the link function, outputting the rate of success (modeling the variant allele frequency), with the linear mixed model in Eq. (1) operating on its logits.

2. *Gaussian:* This approach abstracts the discrete nature of genotypes by employing a Gaussian distribution. In this context, the model simplifies to:

$$\boldsymbol{g} \sim \mathcal{N}\left(\boldsymbol{F}\boldsymbol{\alpha}, \sigma_X^2 \mathcal{K}(\boldsymbol{X}) + \sigma_n^2 \boldsymbol{I}_N\right), \tag{2}$$

   which has been extensively applied in multivariate genetic analyses [15, 10, 9, 16]. While in previous work phenotypes were the outcome variables and multiple genetic variants were inputs to a simple covariance function, we here consider a single genetic variant as the outcome variable, with the multivariate phenotype serving as the input to the covariance function.

As our experiments confirmed sufficient calibration and power of the Gaussian likelihood approach, we opted for it in all experiments.

**Choice of Kernel.** The covariance function $\mathcal{K}(\boldsymbol{X})$ produces an $N \times N$ covariance matrix that characterizes the relationships between individuals based on the histological embeddings. A commonly used function is the linear covariance: $\mathcal{K}_{\text{linear}}(\boldsymbol{X}) = \boldsymbol{X}\boldsymbol{X}^T$ [15, 10, 9, 16], which models linear effects of the embeddings $\boldsymbol{X}$, similar to Bayesian linear regression [2]. However, we observed that using a cosine similarity function provided better calibration of P values [14], ensuring unit diagonal elements of the output covariance. This covariance can be expressed as a linear covariance of transformed features $\tilde{\boldsymbol{X}}$, specifically, $\mathcal{K}_{\text{linear}}(\boldsymbol{X}) = \tilde{\boldsymbol{X}}\tilde{\boldsymbol{X}}^T$, where each row of $\tilde{\boldsymbol{X}}$ is obtained by normalizing the corresponding row of $\boldsymbol{X}$ to have an $L^2$ norm of 1.

**Score Test.** To test for association between a single genetic variant and histological embeddings, we assessed $\sigma^2_{\boldsymbol{X}} > 0$ in the model (Eq. (1)) using a score test [15, 9, 16]. For the Gaussian model, the test statistic is given by:

$$Q = \frac{1}{2}\boldsymbol{g}^T \boldsymbol{P} \mathcal{K}(\boldsymbol{X}) \boldsymbol{P} \boldsymbol{g}, \tag{3}$$

where:

$$\boldsymbol{P} = \frac{1}{\hat{\sigma^2_n}} \left( \boldsymbol{I} - \boldsymbol{F} \left( \boldsymbol{F}^T \boldsymbol{F} \right)^{-1} \boldsymbol{F}^T \right), \tag{4}$$

and $\hat{\sigma^2_n}$ is the maximum likelihood estimator of $\sigma^2_n$ under the null model in Eq. (2) (with $\sigma^2_{\boldsymbol{X}} = 0$). Asymptotically, the test statistic $Q$ follows a mixture of $\chi^2$ distributions under the null hypothesis:

$$Q \sim \sum_i \phi_i \chi^2_1, \tag{5}$$

where:

$$\boldsymbol{\phi} = \text{eigenvalues} \left( \frac{1}{2} \boldsymbol{P}^{\frac{T}{2}} \mathcal{K}(\boldsymbol{X}) \boldsymbol{P}^{\frac{1}{2}} \right). \tag{6}$$

Full details of this derivation can be found in [9]. P values are obtained using the Davies method [4]. Following [15], we use Liu saddlepoint approximation [11] to obtain P values when the Davies method fails to converge.

**Efficient Implementation.** To ensure that HistoGWAS is scalable to large cohorts, we exploit the fact that the dimensionality of the embeddings is typically much lower than the number of individuals ($L \ll N$) [13]. This allows us to achieve linear scaling with respect to the number of individuals. The test statistic $Q$ is computed as follows:

$$\hat{\boldsymbol{\alpha}} = \left( \boldsymbol{F}^T \boldsymbol{F} \right)^{-1} \boldsymbol{F}^T \boldsymbol{g}, \tag{7}$$

$$\hat{\boldsymbol{y}} = \left( \boldsymbol{g} - \boldsymbol{F} \hat{\boldsymbol{\alpha}} \right), \tag{8}$$

$$Q = \frac{1}{2\hat{\sigma^2_n}} \left( \tilde{\boldsymbol{X}}^T \hat{\boldsymbol{y}} \right)^T \left( \tilde{\boldsymbol{X}}^T \hat{\boldsymbol{y}} \right). \tag{9}$$

Using the cyclic property of eigenvalues, the non-zero eigenvalues of $\frac{1}{2}\boldsymbol{P}^{\frac{T}{2}} \tilde{\boldsymbol{X}} \tilde{\boldsymbol{X}}^T \boldsymbol{P}^{\frac{1}{2}}$ can be obtained as the eigenvalues of the matrix $\boldsymbol{\Lambda} = \frac{1}{2} \tilde{\boldsymbol{X}}^T \boldsymbol{P} \tilde{\boldsymbol{X}}$, which is an $L \times L$ matrix. This matrix can be efficiently computed as follows:

$$\widehat{\boldsymbol{X}} = \boldsymbol{P} \tilde{\boldsymbol{X}} = \tilde{\boldsymbol{X}} - \boldsymbol{F} \left( \boldsymbol{F}^T \boldsymbol{F} \right)^{-1} \boldsymbol{F}^T \tilde{\boldsymbol{X}}, \tag{10}$$

$$\boldsymbol{\Lambda} = \frac{1}{2} \tilde{\boldsymbol{X}}^T \widehat{\boldsymbol{X}}. \tag{11}$$

All operations required to compute $Q$ and $\boldsymbol{\Lambda}$ scale linearly with $N$. Furthermore, both the computation of the eigenvalues of $\boldsymbol{\Lambda}$ and the calculation of P values depend on the dimensionality of the embeddings, $L$, rather than on $N$. The eigenvalue computation scales cubically with $L$.

#### 1.4 Visualization of Genetic Effects on Histology

To illustrate histological alterations linked to significant genetic variants, we describe the main steps involving latent space mixed models, embedding interpolation, and semantic decoding:

1. **Fit Linear Mixed Model:** We fit a linear mixed model where the vector $\boldsymbol{g}$ ($N \times 1$) of genotype values for a specific variant is modeled as the outcome, influenced by covariates $\boldsymbol{F}$ ($N \times K$) with fixed effects $\boldsymbol{\alpha}$ ($K \times 1$), and individual-level embeddings $\boldsymbol{X}$ ($N \times L$) with random effects $\boldsymbol{\beta}$ ($L \times 1$). Covariates include an intercept, sex, age, the leading four genetic principal components, and one-hot encoded variables for the type of death.

2. **Define Genetic Effect Axis:** After fitting the linear mixed model, the mean of the posterior distribution of the random effect $\boldsymbol{\beta}$ is given by:

$$\widehat{\boldsymbol{\beta}} = \widehat{\sigma^2_{\boldsymbol{X}}} \boldsymbol{X}^T \left( \widehat{\sigma^2_{\boldsymbol{X}}} \boldsymbol{X}\boldsymbol{X}^T + \widehat{\sigma^2_n} \boldsymbol{I}_N \right)^{-1} (\boldsymbol{g} - \boldsymbol{F}\widehat{\boldsymbol{\alpha}}), \tag{12}$$

   where $\widehat{\cdot}$ denotes the maximum likelihood estimator. The vector $\widehat{\boldsymbol{\beta}}$ is an $L \times 1$ vector that defines the phenotypic direction most predictive of the genotype. We name this direction the genetic effect axis, leveraging linear projections of embeddings along this axis to quantify and visualize the affected phenotype.

3. **Projection of Individual-Level Embeddings:** The projection of individual-level embeddings onto the genetic effect axis is obtained through the best linear unbiased predictor (BLUP):

$$\boldsymbol{g}_{\text{BLUP}} = \boldsymbol{X}\widehat{\boldsymbol{\beta}} = \underbrace{\widehat{\sigma^2_{\boldsymbol{X}}} \boldsymbol{X}\boldsymbol{X}^T \left( \widehat{\sigma^2_{\boldsymbol{X}}} \boldsymbol{X}\boldsymbol{X}^T + \widehat{\sigma^2_n} \boldsymbol{I}_N \right)^{-1}}_{\boldsymbol{H}} \underbrace{(\boldsymbol{g} - \boldsymbol{F}\widehat{\boldsymbol{\alpha}})}_{\boldsymbol{g}_{\text{R}}}, \tag{13}$$

   where we introduced the matrix $\boldsymbol{H}$ and residuals $\boldsymbol{g}_{\text{R}}$. To avoid overfitting from this in-sample estimator, we use the leave-one-out (LOO) estimator proposed in [12]:

$$\boldsymbol{g}_{\text{LOO},i} = \frac{\boldsymbol{g}_{\text{BLUP},i} - \boldsymbol{H}_{i,i}\boldsymbol{g}_{\text{R},i}}{1 - \boldsymbol{H}_{i,i}}, \tag{14}$$

   where $\boldsymbol{g}_{\text{BLUP}}$ is the in-sample predictor, and $\boldsymbol{H}_{i,i}$ represents the $i$-th diagonal element of the projection matrix. This leave-one-out estimator is used to compute individual-level genotype axis scores for visualization in **Figure 4** and Supplementary Figures 8.

4. **Projection of Patch-Level Embeddings:** Since the model is linear in the slide-level embeddings and these embeddings are obtained as a linear average of the patch-level embeddings (via average pooling), we can apply the same projection to all patches to obtain patch-level phenotypic scores $\boldsymbol{s}$:

$$\boldsymbol{s} = \boldsymbol{X}_{\text{patch}}\widehat{\boldsymbol{\beta}}, \tag{15}$$

   where $\boldsymbol{X}_{\text{patch}}$ indicates the #patches $\times$ $L$ matrix of patch-level embeddings.

5. **Interpolate Between Embeddings:** First, we compute extreme embeddings for interpolation as the average of patches at both extremes of the distribution

of $\boldsymbol{s}$. Specifically, we average all patches in the 1st-5th percentiles of $\boldsymbol{s}$ to obtain $\boldsymbol{z}_m$, and all patches in the 95th-99th percentiles of $\boldsymbol{s}$ to obtain $\boldsymbol{z}_M$. Next, we linearly interpolate between $\boldsymbol{z}_m$ and $\boldsymbol{z}_M$ as:

$$\boldsymbol{z}(\alpha) = (1-\alpha)\boldsymbol{z}_m + \alpha\boldsymbol{z}_M, \tag{16}$$

where $\alpha \in [0, 1]$.

6. **Decode Interpolated Embeddings:** Each interpolated embedding $\boldsymbol{z}(\alpha)$ is decoded using the semantic decoder. Given the stochastic nature of our decoder, multiple visual interpretations can be generated by varying the input noise, allowing for several visualizations of the continuum of histological changes (**Supplementary Figures 11**).

All calculations are linear in the cohort size due to leveraging the fact that the embedding dimension is much smaller than the number of individuals. This is achieved by factorizing all operations appropriately and utilizing the Woodbury identity and matrix determinant lemma to solve linear systems, compute inverses, and determine log determinants [3, 2, 13]. Moreover, fitting the linear mixed model uses the $\delta$ reparameterization introduced in [8], where fast grid search on $\delta$ is achieved leveraging that all MLEs can be computed in closed form for any fixed value of $\delta$. All optimized models are made available.

#### 1.5 Power Analysis for HistoGWAS

We simulated 64-dimensional individual-level embeddings as a sum of contributions from covariates, a single genetic variant, and Gaussian noise. Each component was simulated as follows:

1. **Covariate Effects:** We generate covariates $\boldsymbol{F}$ (an $N \times K$ matrix) to capture effects from sex (modeled as a 50/50 Bernoulli distribution), age (uniform distribution between 40 and 80), and the four genetic principal components (random normal distribution). The corresponding effects matrix $\boldsymbol{\alpha}$ ($K \times L$) is sampled from a random normal distribution. The contribution of covariates is computed as:

$$\boldsymbol{X}_c = \boldsymbol{F} \cdot \boldsymbol{\alpha}, \tag{17}$$

and normalized to have an average variance of $v_c$ across dimensions (we set $v_c = 20\%$):

$$\boldsymbol{X}_c = \sqrt{\frac{v_c}{\text{mean}(\text{var}(\boldsymbol{X}_c, 0))}}\boldsymbol{X}_c. \tag{18}$$

2. **Genetic Effects:** We generate a genotype vector $\boldsymbol{g}$ ($N \times 1$) from a binomial distribution with two trials and minor allele frequencies uniformly distributed between 5% and 20%. The genetic effects matrix $\boldsymbol{\beta}$ ($1 \times L$) is sampled from an iid random normal distribution. The contribution of the genetic variant is computed as:

$$\boldsymbol{X}_g = \boldsymbol{g} \cdot \boldsymbol{\beta}, \tag{19}$$

and normalized to have an average variance of $v_g$ across dimensions:

$$\boldsymbol{X}_g = \sqrt{\frac{v_g}{\text{mean}(\text{var}(\boldsymbol{X}_g, 0))}}\boldsymbol{X}_g. \tag{20}$$

3. **Gaussian Noise:** Gaussian noise $\boldsymbol{X}_n$ $(N \times L)$ is generated as:

$$\boldsymbol{X}_n = \text{np.random.randn}(N, L), \tag{21}$$

and normalized such that the average variance of $\boldsymbol{X}_c + \boldsymbol{X}_g + \boldsymbol{X}_n$ across dimensions is approximately 1:

$$\boldsymbol{X}_n = \sqrt{\frac{1 - v_c - v_g}{\text{mean}(\text{var}(\boldsymbol{X}_n, 0))}} \boldsymbol{X}_n. \tag{22}$$

### 2 Supplementary Datasets

**Supplementary Dataset 1: Overview of Genotype and Histology Data Across Eleven Tissue Types from the GTEx Project.** This table summarizes the comprehensive dataset derived from the GTEx project, capturing the intersection of genetic and histological analyses across eleven selected tissue types. It presents the number of individuals with genotype data, expression data, and the crucial intersection of histological imaging with both genetic and expression information. The progression from the initial number of extracted histological patches to those retained after quality processing, and finally to patches classified into distinct histological cluster signatures, is detailed. The tissues are ranked based on the count of individuals for whom both histological images and genotype data are available, emphasizing the depth of integrated data for each tissue type.

**Supplementary Dataset 2 : Gene Expression Associations with Histological Cluster Signatures Across Tissues.** A comprehensive multi-tab Excel file presenting the results from the association analysis linking histological cluster signature abundance with gene expression levels across various tissues. Each tab provides association statistics for a single cluster signature. This dataset can be used to further characterize cluster signatures, offering a rich resource for further biological exploration of our embeddings.

**Supplementary Dataset 3 : Results from Post-GWAS Analyses of Genome-Wide Significant Loci.** Organized in a structured multi-tab Excel file, this dataset offers a concise summary of post-GWAS analyses, including info on all genome-wide significant variants (1 tab), association statistics of each lead locus with expression levels of highly variable genes in the respective tissue (4 tabs, one for each of 4 variants), and results from pathway enrichment analysis in both upregulated (4 tabs, one for each of 4 variants) and downregulated gene sets (4 tabs, one for each of 4 variants).

#### 3 Supplementary Figures

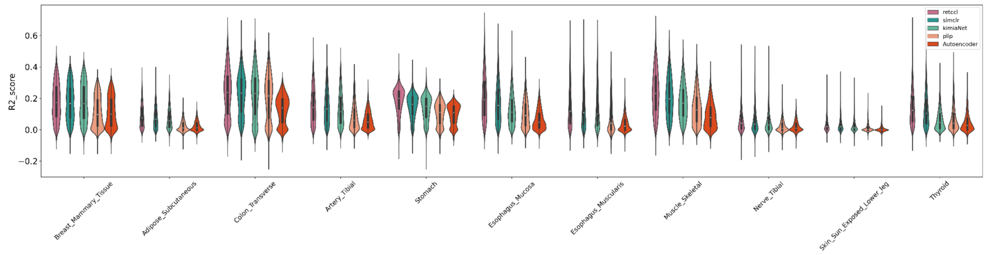

**Supplementary Figure 1: Comparative Analysis of Encoding Models for Gene Expression Prediction.** Shown is the violin plot showing the cross-gene distribution of $R^2$ between observed and histologically predicted gene expression levels for test set individuals across the 11 analyzed tissues. This metric quantifies the predictive accuracy of gene expression from individual-level histological embeddings obtained from different models: RetCCL, SimCLR, KimiaNet, PLIP and Autoencoder (**Methods**). Notably, the RetCCL contrastive learning model exhibits the highest predictive performance across tissues and was selected as HistoGWAS encoder.

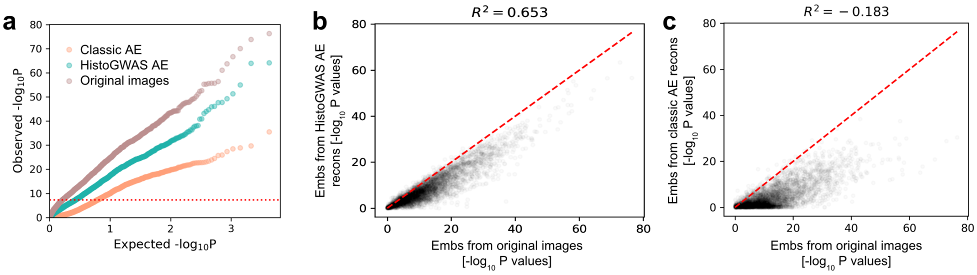

**Supplementary Figure 2: Evaluation of Semantic Autoencoder through Expression Prediction from Reconstructed Images in Thyroid tissue.** (**a**) QQ plots of P values of association between predicted and observed gene expression levels on test individuals, using embeddings of original images, embeddings of reconstructions from a classic autoencoder (classic AE), or embeddings of reconstructions from HistoGWAS autoencoder (HistoGWAS AE, Methods). This evaluation was performed exclusively in thyroid tissue, with the predictive model consistently trained using embeddings of original images on the training set for all comparisons. (**b**) Scatter plot analysis of gene expression prediction statistics (-log10 P) employing embeddings of reconstructions from HistoGWAS AE (y-axis) versus embeddings of original images (x-axis) across genes in thyroid tissue. The coefficient of determination ($R^2$) is also reported to illustrate the predictive performance. (**c**) Analogous to (**b**), but comparing prediction statistics using embeddings of reconstructions from classic AE (y-axis) versus embeddings of original images (x-axis) across genes in thyroid tissue.

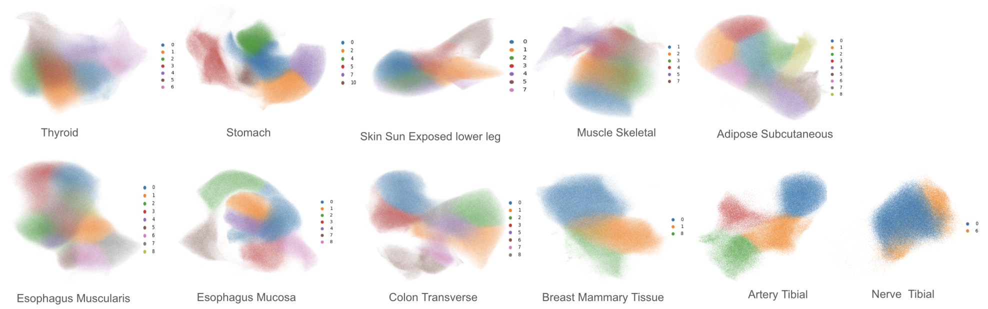

**Supplementary Figure 3: Unsupervised Data Analysis Reveals Cluster Signatures Across Tissues.** This figure illustrates the results of an unsupervised data analysis, showcasing 64 unique cluster signatures identified across eleven tissues. Shown is the UMAP (Uniform Manifold Approximation and Projection) obtained for each tissue, with colors delineating the distinct cluster signatures as identified through Leiden clustering.

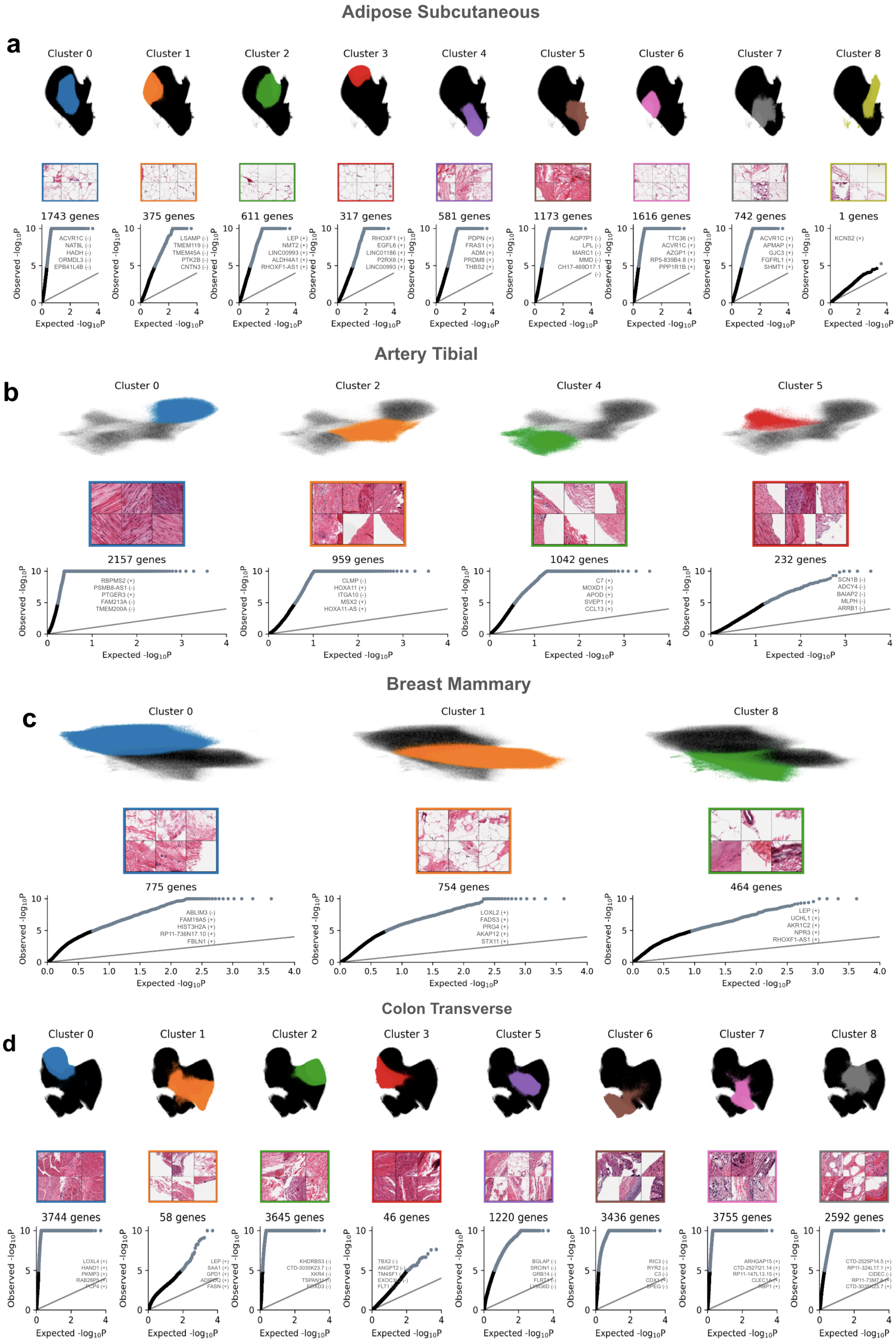

**Supplementary Figure 4: Validation of Cluster Signatures across Multiple Tissues via Gene Expression Correlation.** *Continued on the next page.*

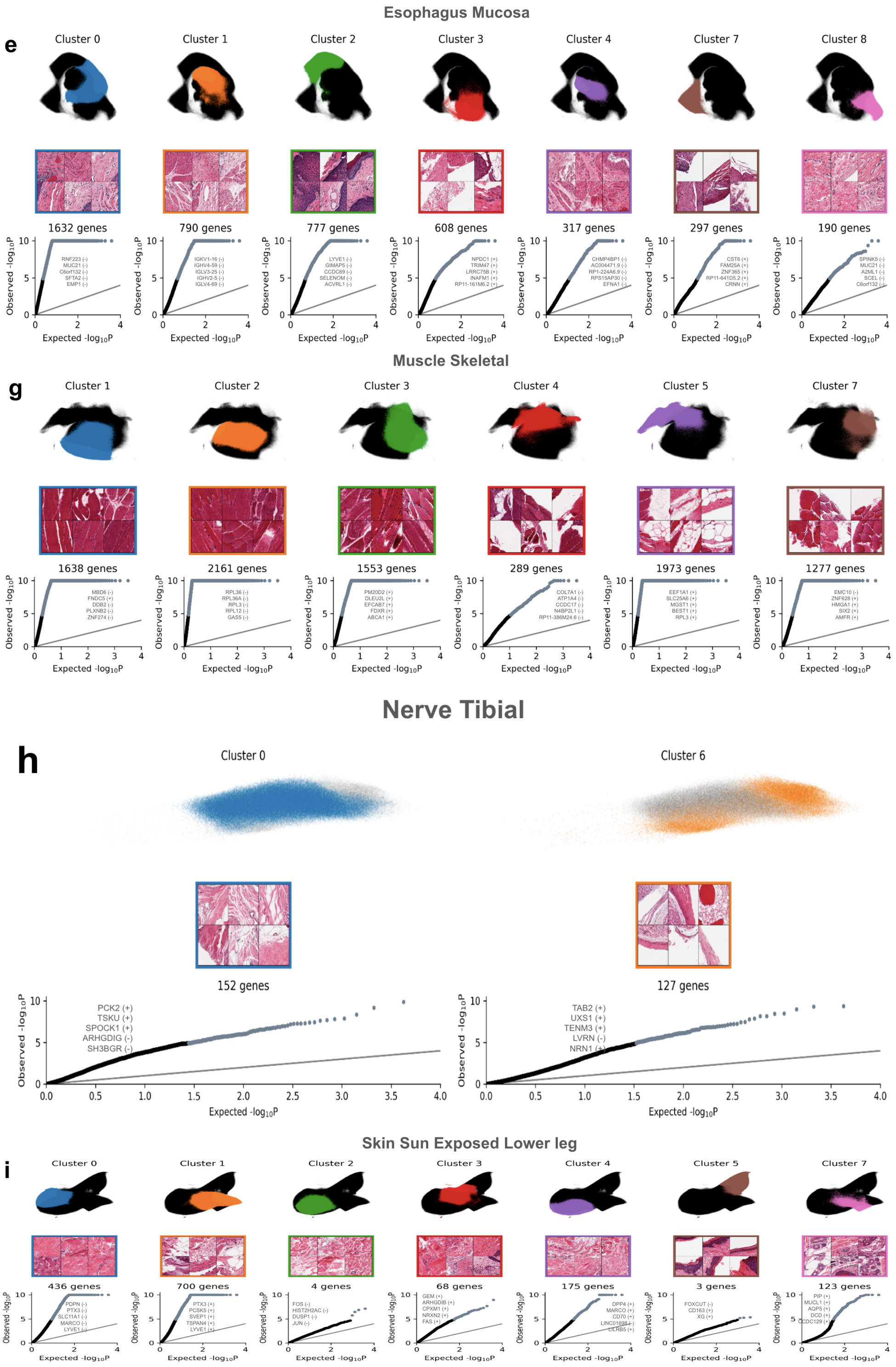

**Supplementary Figure 4: Validation of Cluster Signatures across Multiple Tissues via Gene Expression Correlation.** *Continued on the next page.*

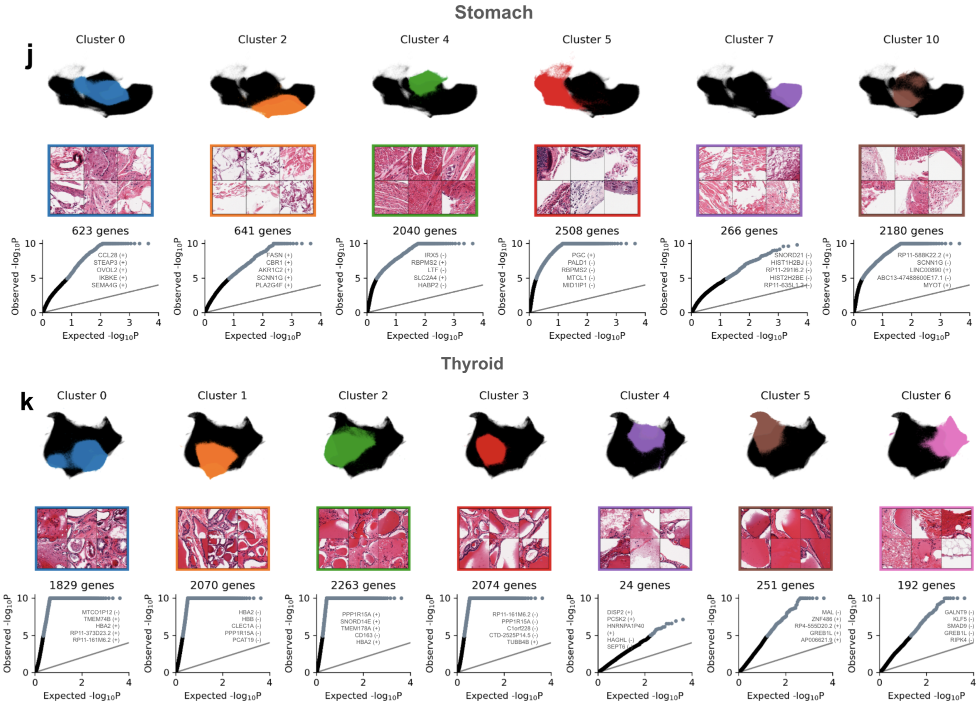

**Supplementary Figure 4: Validation of Cluster Signatures across Multiple Tissues via Gene Expression Correlation.** This figure illustrates the correlation between the fraction of patches associated with distinct cluster signatures within each slide and their subsequent association with gene expression levels in the corresponding tissue. Shown are cluster 64 cluster signatures across the 11 analyzed tissues: Adipose Subcutaneous (**a**), Artery Tibial (**b**), Breast Mammary Tissue (**c**), Colon Transverse (**d**), Esophagus Mucosa (**e**), Esophagus Muscularis (**f**), Muscle Skeletal (**g**), Nerve Tibial (**h**), Skin Sun Exposed Lower Leg (**i**), Stomach (**j**), and Thyroid (**k**). Uniform Manifold Approximation and Projection (UMAP) visualizations display the distribution of cluster signatures, accompanied by exemplar patches for each tissue type. Quantile-Quantile (QQ) plots of P values underscore the association between cluster signature abundance within slides and gene expression, with the top five genes for each tissue marked, showing the directionality of their expression changes (positive associations marked with (+) and negative with (-)), reflecting over or underexpression correlated with cluster signature prevalence. Full results from expression correlation analysis can be found in Supplementary Dataset 2.

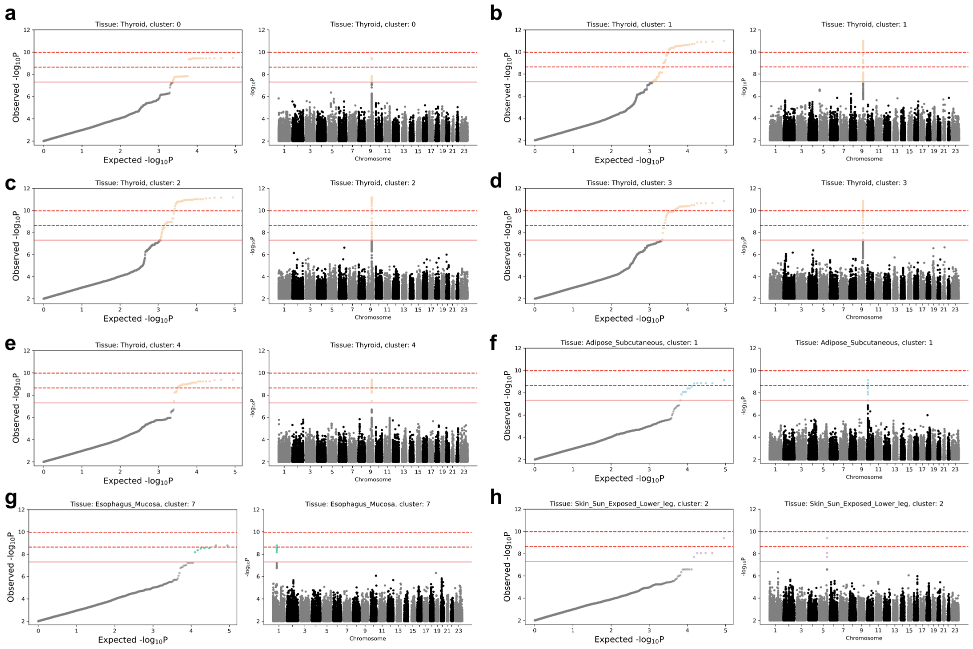

**Supplementary Figure 5: Manhattan and QQ Plots for cluster signatures with Genome-Wide Significant Loci.** Shown are QQ plots (left) and Manhattan plots (right) for cluster signatures with genome-wide significant loci, highlighting five signature clusters in thyroid tissue associated with rs7030256 (**a-e**), one in adipose subcutaneous associated with variant rs1432621 (**f**), one in esophagus mucosa associated with variant rs3766325 (**g**), and one in sun-exposed skin associated with rs1432621 (**h**). We show three P value thresholds for statistical significance: the standard significance level, P < 5.10-8, $P < 323 \cdot 10 - 9$ (corresponding to FWER < 20%, computed through permutations, (**Methods**)), and P ¡ 1.5310-10 (corresponding to FWER 5%, computed through permutations).

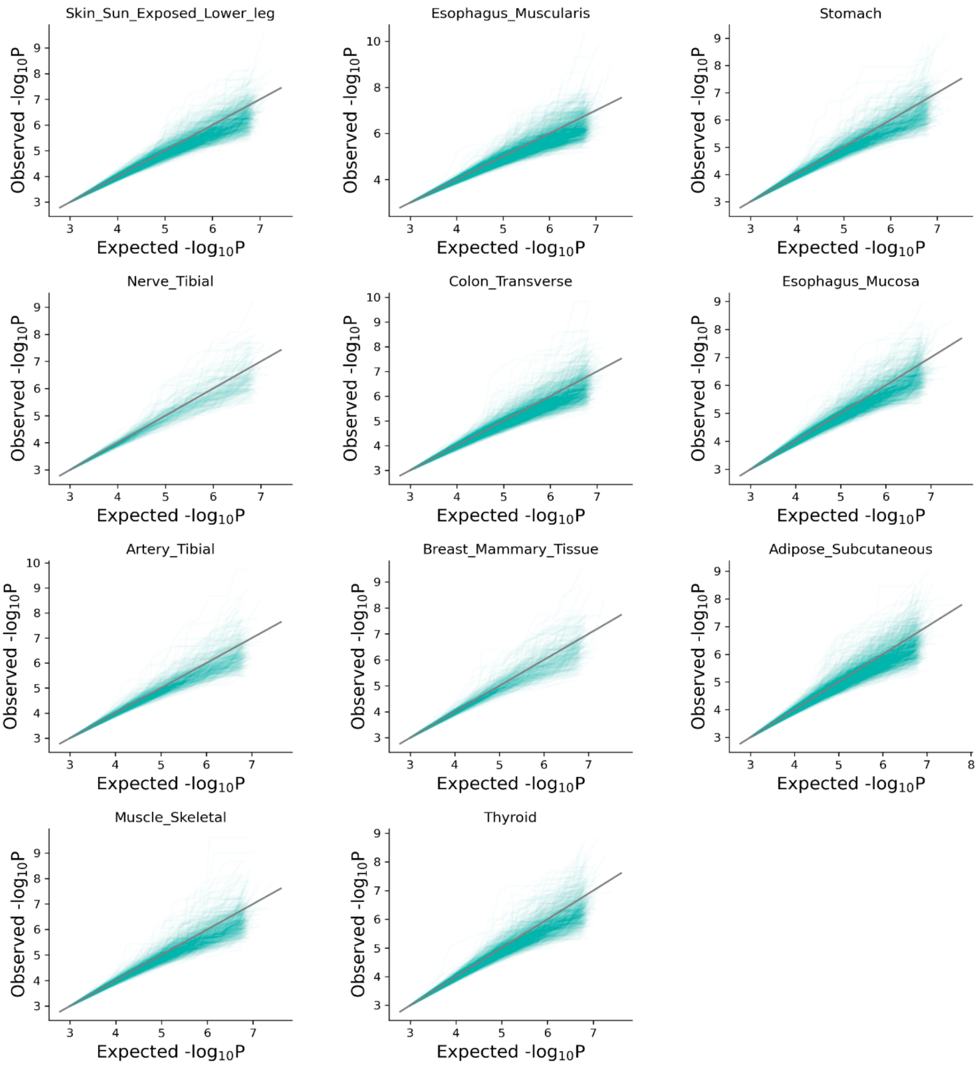

**Supplementary Figure 6: Calibration of P-values from HistoGWAS Under Permuted Data** Displayed are QQ plots of P-values obtained from genome-wide analysis using HistoGWAS for 68 cluster signatures, involving approximately 5 million genetic variants, under permuted data conditions (**Methods**). Each panel illustrates QQ plots across cluster signatures for 100 permutations within a specific tissue.

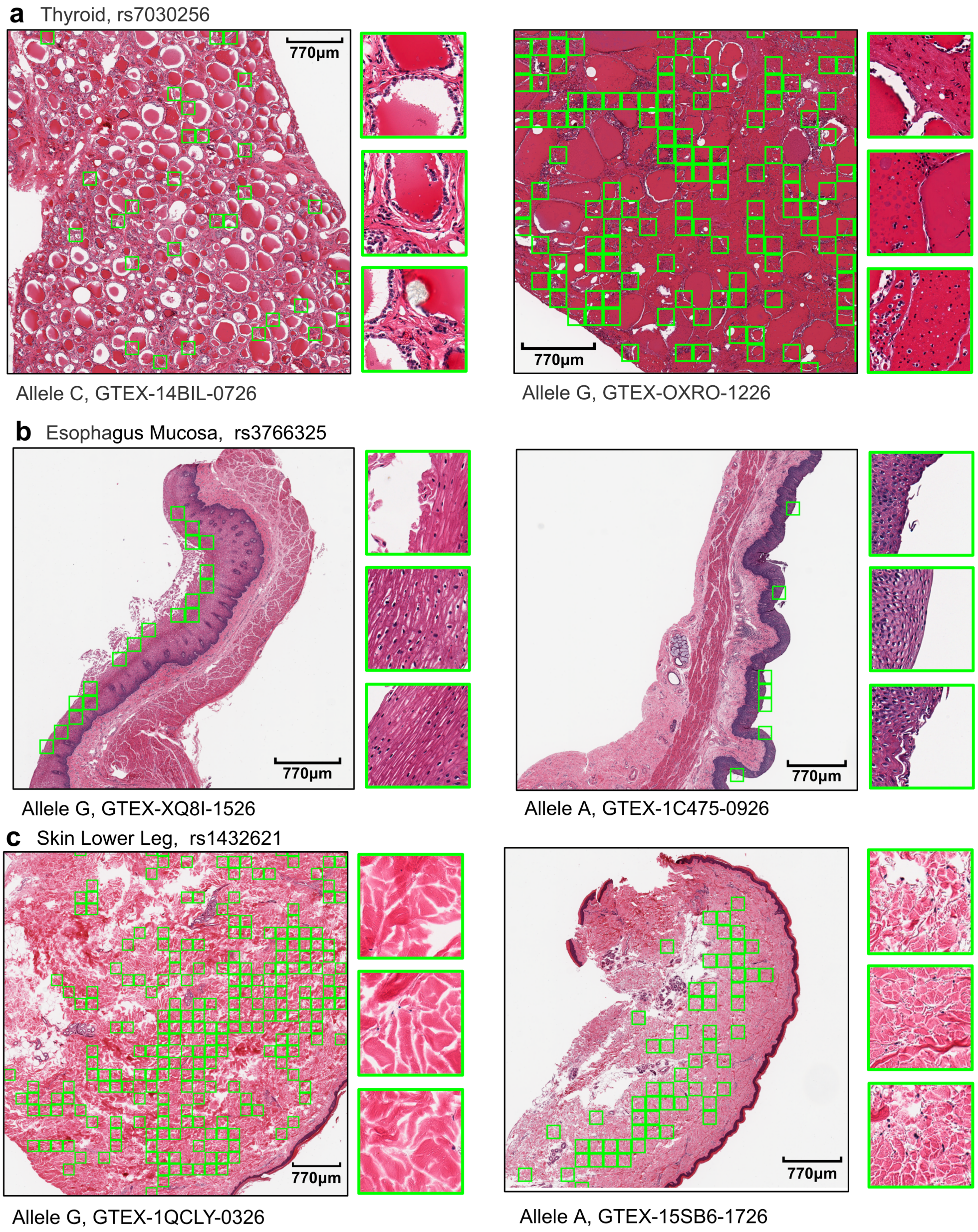

**Supplementary Figure 7: Genotype-Driven Histological Phenotypes Visualized on Whole-Slide Images.** Continued on the next page.

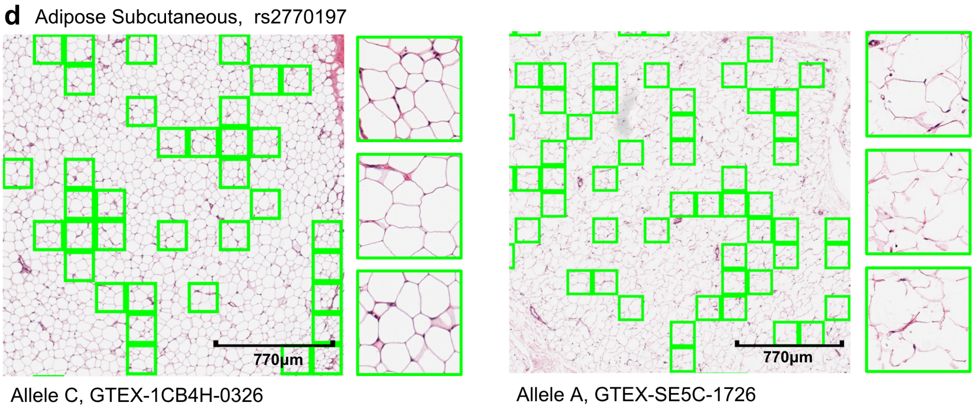

**Supplementary Figure 7: Genotype-Driven Histological Phenotypes Visualized on Whole-Slide Images.** Shown are sections of whole-slide images in which we have identified and highlighted tiles that demonstrate a close association with specific alleles at significant genetic loci (refer to Methods). For every allele, 3 tiles are magnified to provide a detailed view. This approach aims to visually display the histological variations attributable to different genetic variants within the tissue context. The analysis encompasses tissue samples from Thyroid (a), Esophagus Mucosa (b), Skin Lower Leg (c) and Adipose Subcutaneous (d). A detailed visualization of each GTEX slide is accessible at `https://gtexportal.org/home/histologyPage` using the respective tissue sample IDs.

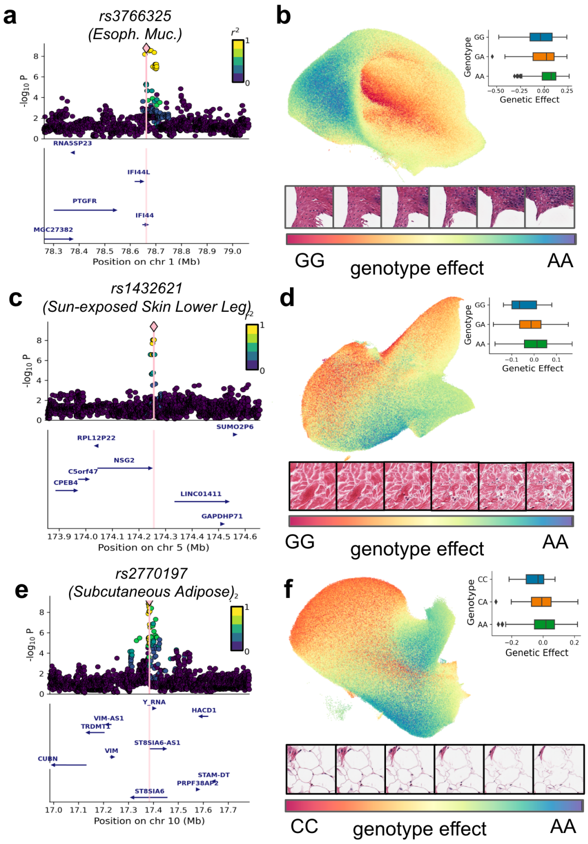

**Supplementary Figure 8: Visualization of Genomic Context and Histological Impact of selected Genetic Variants.** Analogous to Figure 4, this figure presents the locus zoom plot and histological effects of the remaining three significant variants, rs3766325 in Esophagus Mucosa **a-b**, rs1432621 in Skin Lower Leg **c-d** and rs2770197 in Subcutaneous Adipose tissue **e-f**.

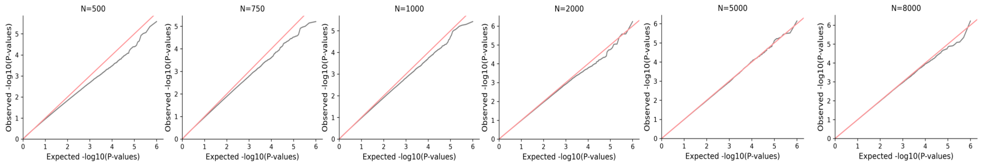

**Supplementary Figure 9: Assessment of calibration of HistoGWAS in Simulated Datasets with no Genetic Effects.** QQ plots illustrating the calibration of P values from HistoGWAS under null conditions (no genetic effects) across various simulated cohort sizes **Methods**.

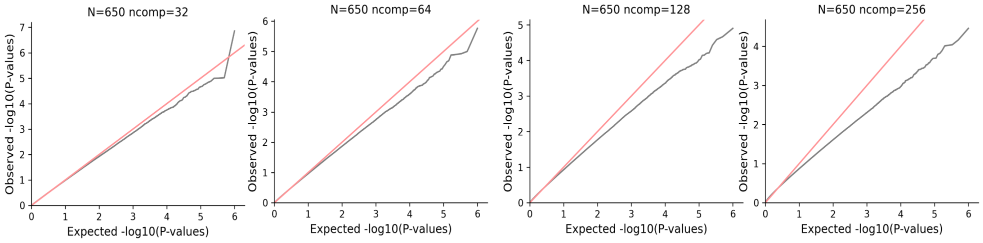

**Supplementary Figure 10: Calibration of P-values in Simulated Datasets with No Genetic Effects Across Different Embedding Dimensions.** This figure presents QQ plots of P-values derived from HistoGWAS applied to simulated datasets under a null model (no genetic effects), matching the smallest cohort size used in our study (N=650). Each plot varies the number of latent embedding dimensions (ncomps), corresponding to the number of principal components of embeddings **Methods**. The analysis reveals an increasing deflation of P-values with an increasing number of components, which informed our decision to utilize 64 components in our genetic analyses.

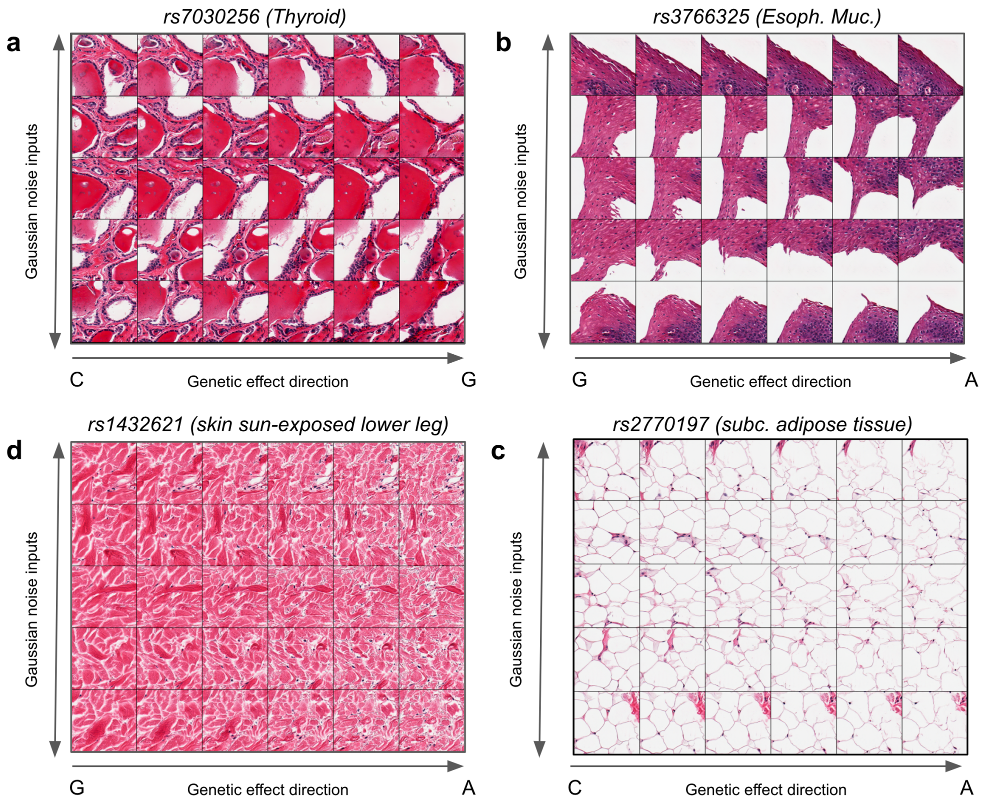

**Supplementary Figure 11: Visualization of Histological Variations Induced by Genetic Variants Across Different Decoder Input Noises.** This figure demonstrates how varying samples of Gaussian noise input to the semantic decoder influence the visualization of histological changes associated with the four detected genome-wide significant loci. Each panel displays a range of histological outcomes that reflect the stochastic nature of the decoding process, enabling thorough evaluation **Methods**.

[2] Francesco Paolo Casale. Multivariate linear mixed models for statistical genetics. PhD thesis, 2016.

[14] Hinrich Schütze, Christopher D Manning, and Prabhakar Raghavan. Introduction to information retrieval, volume 39. Cambridge University Press Cambridge, 2008.

[15] Michael C Wu, Seunggeun Lee, Tianxi Cai, Yun Li, Michael Boehnke, and Xihong Lin. Rare-variant association testing for sequencing data with the sequence kernel association test. The American Journal of Human Genetics, 89(1):82–93, 2011.

[16] Wei Zhou, Wenjian Bi, Zhangchen Zhao, Kushal K Dey, Karthik A Jagadeesh, Konrad J Karczewski, Mark J Daly, Benjamin M Neale, and Seunggeun Lee. Saige-gene+ improves the efficiency and accuracy of set-based rare variant association tests. Nature genetics, 54(10):1466–1469, 2022.
